# Technical Note: Efficient and accurate estimation of genotype odds ratios in biobank-based unbalanced case-control studies

**DOI:** 10.1101/646018

**Authors:** Rounak Dey, Seunggeun Lee

## Abstract

In genome-wide association studies (GWASs), genotype log-odds ratios (LORs) quantify the effects of the variants on the binary phenotypes, and calculating the genotype LORs for all of the markers is required for several downstream analyses. Calculating genotype LORs at a genome-wide scale is computationally challenging, especially when analyzing large-scale biobank data, which involves performing thousands of GWASs phenome-wide. Since most of the binary phenotypes in biobank-based studies have unbalanced (*case* : *control* = 1 : 10) or often extremely unbalanced (*case* : *control* = 1 : 100) case-control ratios, the existing methods cannot provide a scalable and accurate way to estimate the genotype LORs. The traditional logistic regression provides biased LOR estimates in such situations. Although the Firth bias correction method can provide unbiased LOR estimates, it is not scalable for genome-wide or phenome-wide scale association analyses typically used in biobank-based studies, especially when the number of non-genetic covariates is large. On the other hand, the saddlepoint approximation-based test (fastSPA), which can provide accurate p values and is scalable to analyse large-scale biobank data, does not provide the genotype LOR estimates as it is a score-based test. Here, we propose a scalable method based on score statistics, to accurately estimate the genotype LORs, adjusting for non-genetic covariates. Comparing to the Firth method, our proposed method reduces the computational complexity from *O*(*nK*^2^ + *K*^3^) to *O*(*n*), where *n* is the sample-size, and *K* is the number of non-genetic covariates. Our method is ~ 10x faster than the Firth method when 15 covariates are being adjusted for. Through extensive numerical simulations, we show that the proposed method is both scalable and accurate in estimating the genotype ORs in genome-wide or phenome-wide scale.

## 1 Introduction

Recent developments in genotyping and imputation technologies [Marchini and Howie, 2010, Das et al., 2016], and the availability of electronic health records (EHR)-based phenotypic information in different biobanks [Bycroft et al., 2017, Krokstad et al., 2013, Fritsche et al., 2018], are allowing the researchers to perform Phenome-wide scale genome-wide association studies (GWASs) to investigate the pleiotropic effect [Pendergrass et al., 2013, Hebbring, 2014, Verma et al., 2018] of different variants on multiple phenotypes. Since, subjects in biobanks are usually recruited in a cohort-based design, most binary phenotypes based on biobank data are unbalanced (*case* : *control* = 1 : 10), or often extremely unbalanced (*case* : *control* = 1 : 100).

In case-control GWASs, genotype odds ratios (ORs) quantify the effect of the variants on the phenotypes. Genotype ORs are required for several downstream analyses. For example, in Mendelian randomization studies [Burgess et al., 2013, Bowden et al., 2015, Burgess et al., 2015], they are used to control for unknown confounders to establish causal relationships between different risk factors and biomedical outcomes. Genotype ORs are also used in the inverse variance-weighted meta-analysis method to combine results from multiple studies [Willer et al., 2010, Evangelou and Ioannidis, 2013, Tsoi et al., 2017]. Usually, it is estimated in the logarithmic scale (hence called log-odds ratios or LORs) by maximizing the likelihood of appropriate logistic regression models.

Maximizing the logistic regression likelihood involves updating the estimates iteratively, which requires *O*(*nK*^2^ + *K*^3^) computation per iteration, where *n* is the sample-size, and *K* is the number of covariates to adjust for in the model. Since, the samples in modern large-scale biobank-based GWASs are usually heterogeneous in nature (ancestry, gender, age etc.), the number of covariates, *K* can become substantially large in these studies, and the overall computation time can increase at a fast pace. Moreover, when the case-control ratios are unbalanced and the minor allele counts (MACs) are low, the logistic regression likelihood, and thus the estimated genotype LORs, can be biased [Firth, 1993]. Even though, the maximum penalized likelihood estimation [Ma et al., 2013, Firth, 1993] can provide bias-adjusted LORs, the method is also not scalable to handle biobank-scale datasets. Recently, several score test-based methods were proposed [Dey et al., 2017, Zhou et al., 2017, Dey et al., 2019] to efficiently and accurately test for genotype-phenotype associations for such unbalanced phenotypes. However, as the score tests only fit the model under the null hypothesis of no association, they do not provide any estimate of the genotype LORs.

Here, we propose a fast computation method for genotype LORs in case-control studies using the score statistics of the fastSPA test, which improves the computation time from *O*(*nK*^2^ + *K*^3^) to *O*(*n*). Through applications on simulated data, we demonstrate the performance of our proposed method under different case-control ratios, and for variants with different rarity of alleles.

## 2 Methods

We consider a case-control study with *n* samples, where the phenotype *Y_i_* = 1, 0 denotes the case-control status of the *i*-th subject. Let *G_i_* = 0, 1, 2 denote the MAC for the variant of interest, and *X_i_* denote other covariates or observed confounders, for the *i*-th subject. We denote the outcome vector by *Y* = (*Y*_1_,…, *Y_n_*), the genotype vector by *G* = (*G*_1_,…, *G_n_*), and the covariate matrix by *X* = (*X*_1_…*X_n_*)^⊤^. To model the phenotypes on the genotypes and the covariates, we use the following logistic regression model,

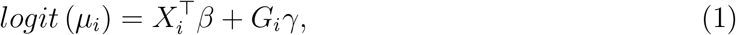

where *μ_i_* = *P*(*Y_i_* = 1|*X_i_, G_i_*), and *β, γ* are the regression parameters. For simplicity of notations, we augmented the intercept term in the covariate matrix *X*. Let *μ* = (*μ*_1_,…, *μ_n_*), and *W* = *diag*(*μ_i_*(1 − *μ_i_*)). Then, the score function and information matrix under model 1 are given by,

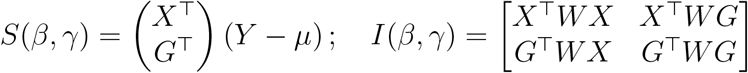

respectively. The standard Newton-Raphson algorithm to obtain the maximum likelihood estimates (mle) 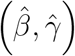 under this model iterates over the following updates,

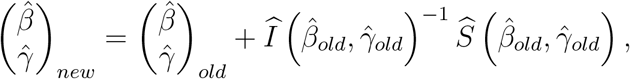

where 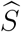 and 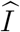 are the estimated score function and the information matrix where the parameters are replaced by their estimates at the current iteration. The computations required to calculate the information matrices and to invert them at each iteration are *O*(*nK*^2^) and *O*(*K*^3^) respectively, where *K* is the number of covariates. When analyzing millions of variants across thousands of phenotypes in models including large numbers of covariates, the computation time can become substantially large.

Our method is motivated by the observations that for relatively small effect sizes in the log-odds scale (LORs), the logistic model behaves closely to a linear model [Pirinen et al., 2013], and the score functions for linear and logistic regressions are algebraically of similar forms. We thus propose to estimate the genotype LORs by fitting a genotype-only model, using analogous estimation techniques used in linear models. For linear regression, the mle for the genotype LOR in the full model 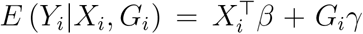 is given by 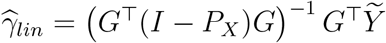, where *P_X_* = *X*(*X*^⊤^*X*)^−1^ *X*^⊤^ is the projection matrix of the covariates, and *Ỹ* = (*I* − *P_X_*)*Y* is the residual from the null model 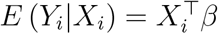. If we use *Ỹ_i_*s as outcomes in the genotype-only model *E*(*Ỹ_i_*|*G_i_*) = *α* + *G_iγ_*, the estimate of *γ* is given by 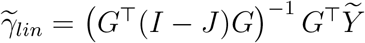, where *nJ* is the *n* × *n* matrix of all elements equal to unity. Therefore, 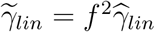, where the scaling factor *f* = {(*G*^⊤^(*I* − *P_X_*)*G*) / (*G*^⊤^(*I* − *J*)*G*)}^1/2^. It is clear from this linear regression setup, that the mles of *γ* from the genotype-only model *E*(*Ỹ_i_*/*f*|*G_i_* = *α* + *fG_iγ_* will be identical to 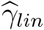. The score function corresponding to this model is,

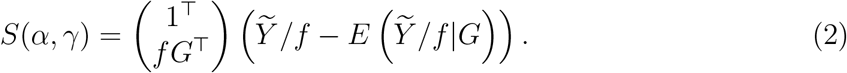

Since, the score functions of the logistic and linear regressions are of the same form, we propose to regress the binary outcomes *Y* on the scaled genotype vector *fG* in a logistic regression, using the same score function as 2. Assuming *G* to be mean-centered, in order to keep the prevalence the same, we use a modified score function,

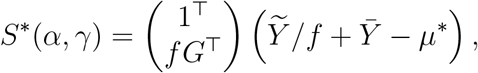

where 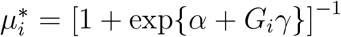. Notice that, this modification does not change the component corresponding to *G* in the score function as *G* is mean-centered. We solve the score equation *S*\*(*α, γ*) = 0 by iterating through the following Newton-Raphson updates,

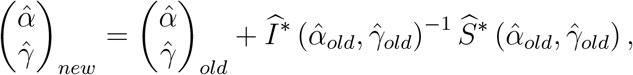

where 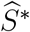 and 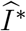 are the estimated score function and the information matrix where the parameters are replaced by their estimates at the current iteration. The information matrix is given by,

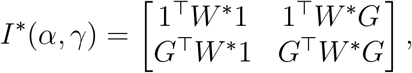

where 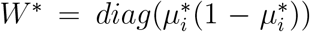. The computations required to calculate the information matrices and to invert them at each iteration are *O*(*n*) and *O*(1) respectively. Therefore, the computation do not depend on *K*, as the information matrices are always of order 2 × 2. Therefore, computationally this method provides substantial improvement over the logistic regression on the full model.

For unbalanced case-control studies, we can further correct the bias in our score function by adjusting it based on the Firth’s bias correction method [Firth, 1993],

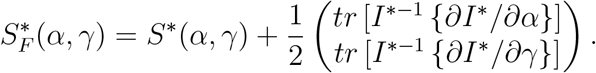

### 2.1 Simulation Studies

To evaluate the proposed method using numerical simulations, we considered three different case-control ratios: balanced (*case* : *control* = 1 : 1), moderately unbalanced (*case* : *control* = 1 : 9), and extremely unbalanced (*case* : *control* = 1 : 99). For each choice of case-control ratio, the phenotypes were simulated based on the following logistic regression model,

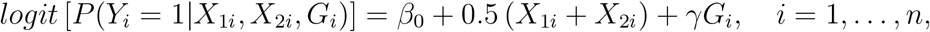

where the two covariates *X*_1*i*_ and *X*_2*i*_ were simulated from *X*_1*i*_ ~ *Bernoulli*(0.5) and *X*_2*i*_ ~ *N*(0, 1), and the intercept *β*_0_ corresponds to the prevalence rate of 1%. We considered two settings to generate the genotypes: (i) uncorrelated with the covariates, where *G_i_* ~ *Binomial*(2, *p*), and (ii) correlated with the covariate *X*_2*i*_s, where *G_i_* ~ *Binomial*(2, *p*\*), *p*\* = *p*(1 − 1/2*r* + *X*_2*i*_/*r*). The minor allele frequencies (MAF) were chosen to be *p* = 0.001, 0.01, 0.05, and the correlation factors (*r*) was chosen to represent allele frequency differences of 0.5*p* (highly correlated) and 0.2*p* (moderately correlated) between the two groups denoted by *X*_2*i*_ = 0 and 1. The uncorrelated setting corresponds to *r* → ∞. Under each case-control ratio, MAF, and *r*, we simulated 200 datasets of sample size *n* = 20000 each using different values of the genotype LOR *γ* (see Figures 8–10). Then, we applied our proposed method (Logistic-Reduced) and Firth’s bias-controlled logistic regression [Ma et al., 2013] under the full model (Logistic-Full), and compared the estimated genotype LORs from these two methods.

To compare the computation times under different numbers of covariates, we simulated datasets with sample size *n* = 20000 from the following logistic regression model,

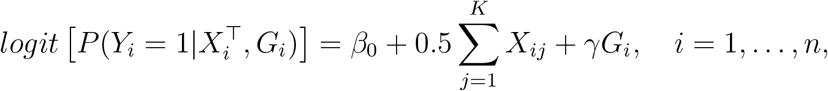

where *K* is the number of covariates. The model is similar to the previously discussed simulation setting, except all the covariates *X_ij_*s follow i.i.d. *N*(0, 1) distribution. The genotype LORs (*γ*) are also selected randomly from *U*(−2, 2) for each dataset. For each of the three case-control ratios, and *K* = 3, 6, 10, 15, 20, 25, we simulated 500 datasets, and compared the computation times for the Logistic-Full and Logistic-Reduced methods.

## 3 Results

We compare the estimated genotype LORs under different case-control ratios and computation times under different numbers of covariates based on extensive numerical simulations.

### 3.1 Simulation Studies

We compare the estimated genotype LORs based on Logistic-Full and Logistic-Reduced methods under different case-control ratios, MAFs, and genotype-covariate correlations. Figures 2,3 and 4 present the scatter plot of the estimated LORs from these two methods for the simulations where genotypes and the covariates are highly correlated, moderately correlated and uncorrelated, respectively. The results show that the estimated LORs are almost identical between these two methods, which suggests that the Logistic-Reduced method provides almost as accurate estimates as the Logistic-Full method. When the case-control ratio is moderately or extremely unbalanced, and the true LOR is very small, there is very little underestimation in Logistic-Reduced compared to Logistic-Full. The comparison of the estimated LORs with the true parameter values for these two methods are presented in Figures 5–10. We notice that, for the rare variants (MAF = 0.01 or 0.001) in the extremely unbalanced case-control setting, both Logistic-Full and Logistic-Reduced methods have extremely large sampling variances, especially when the true LOR is negative. Moreover, the estimates seem to have discrete levels, as the expected number of minor alleles in cases becomes smaller than one. This observation is clearer when MAF = 0.001, where both the Logistic-Full and Logistic-Reduced estimates cluster around zero for negative values of the true LORs.

**Figure 1:**
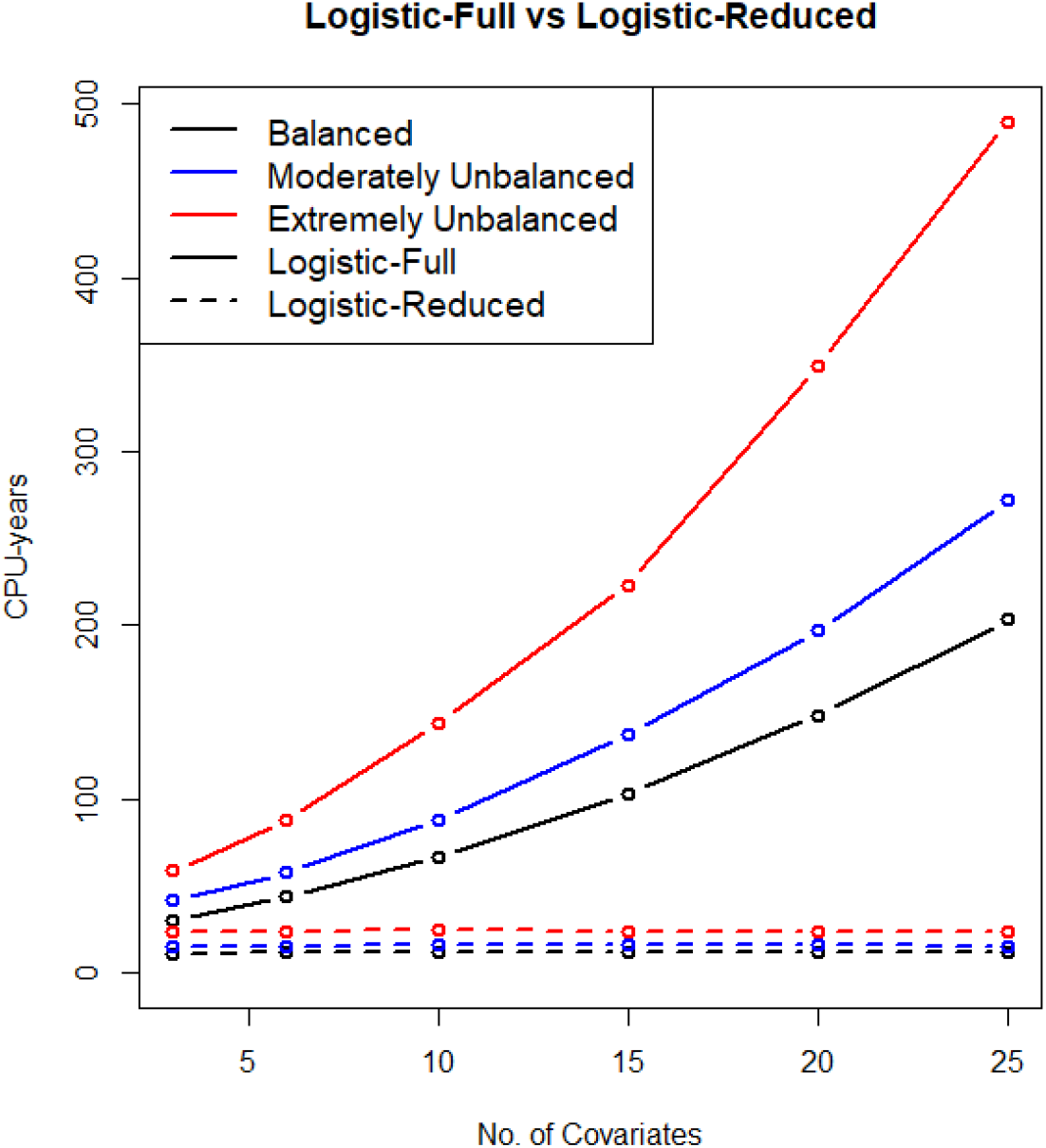
The projected computation times for estimating the genotype LORs using different methods for 10 million variants across 1500 Phenotypes with different number of covariates. The computation times are based on testing 500 simulated variants on an Intel i7 2.70GHz processor and then projecting them onto a PheWAS with 10 million variants and 1500 phenotypes. The solid lines represent the computation times required for Logistic-Full, and the dashed lines represent the computation times required for Logistic-Reduced.

**Figure 2:**
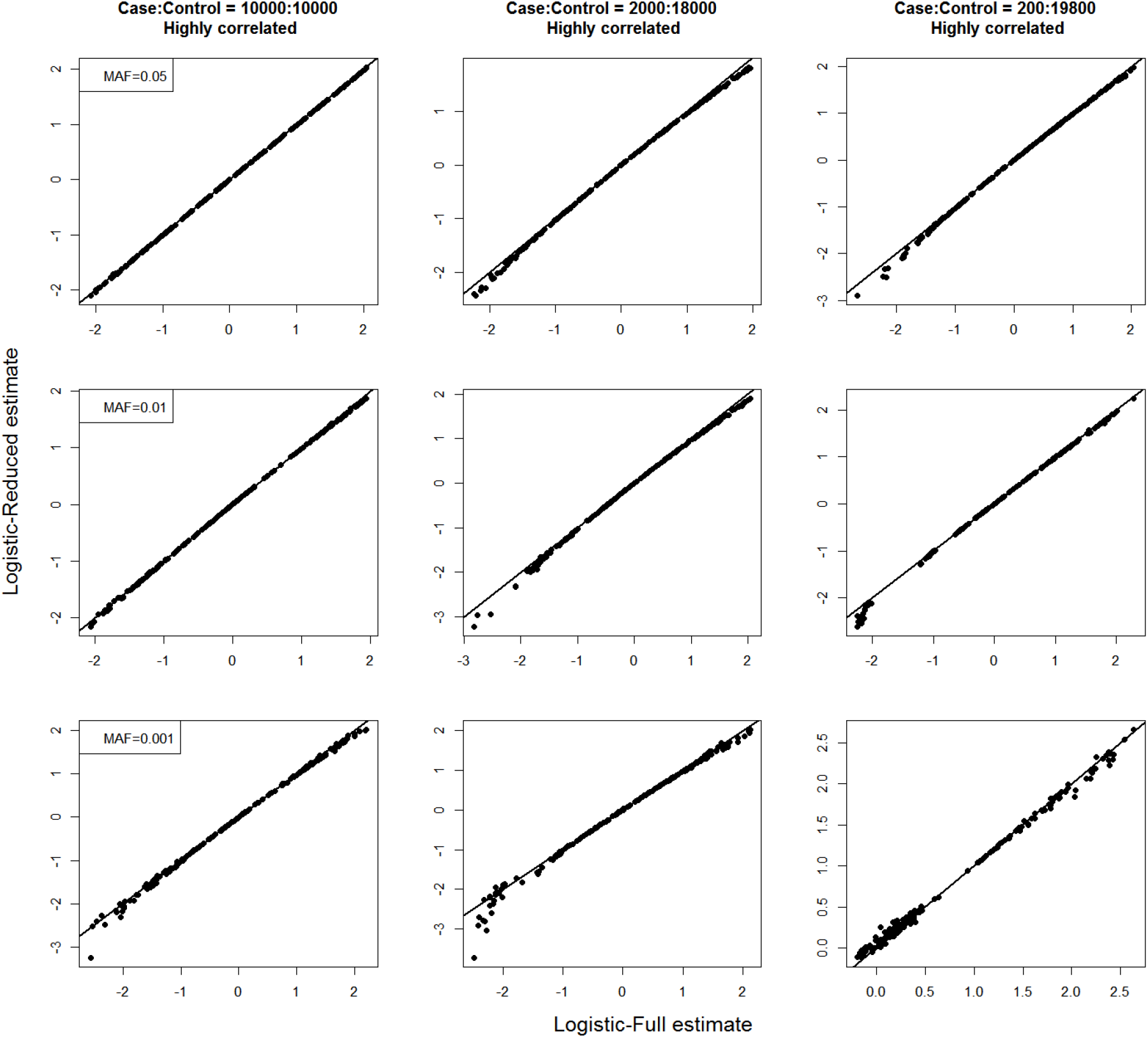
Scatter plots of estimated genotype LORs from Logistic-Full (x-axis) and Logistic-Reduced (y-axis) methods when the genotypes and covariates are highly correlated. From left to right, the panels represent case-control ratios 1 : 1, 1 : 9 and 1 : 99, respectively. From top to bottom, the panels represent MAFs 0.001, 0.01 and 0.05, respectively.

**Figure 3:**
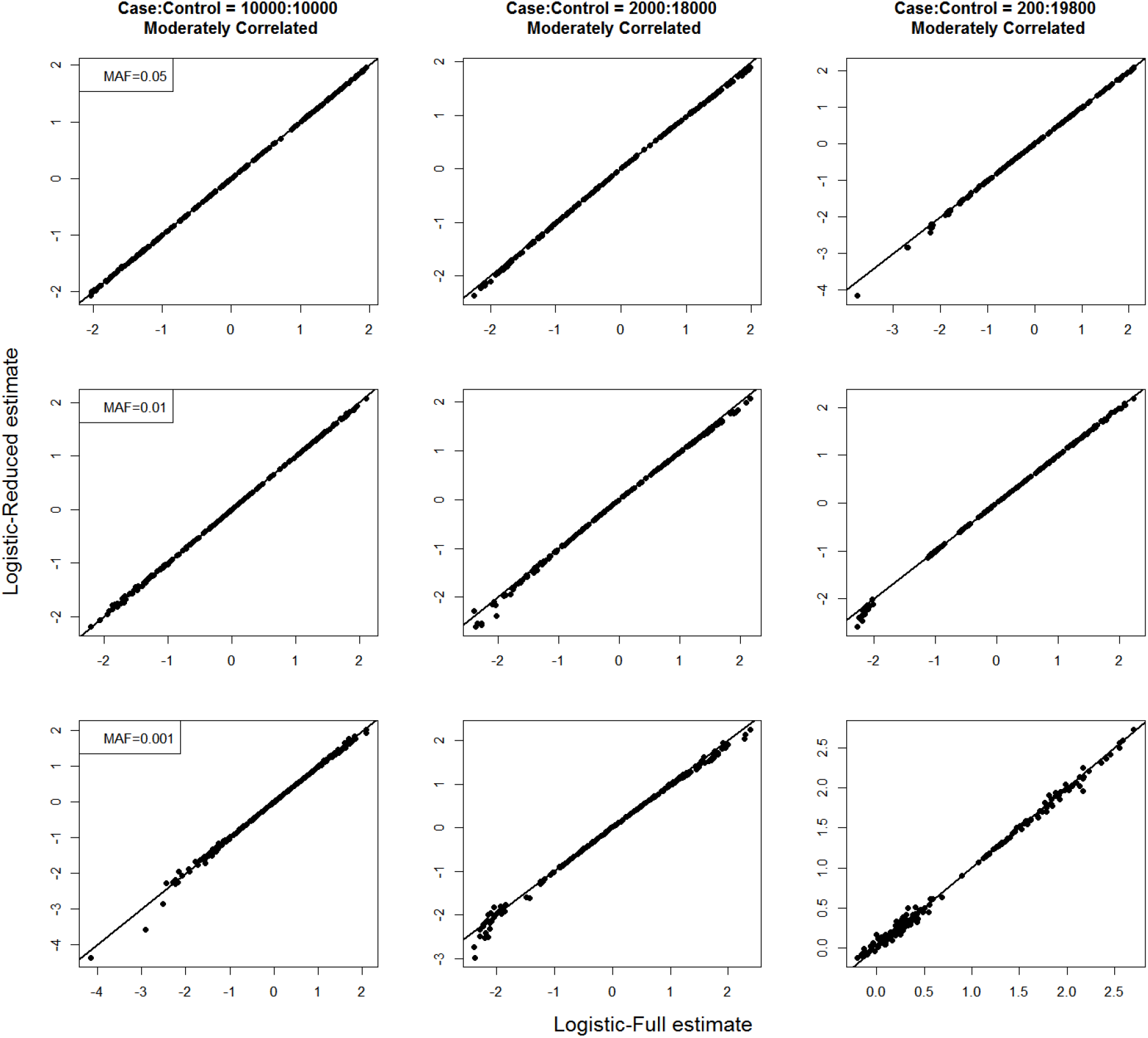
Scatter plots of estimated genotype LORs from Logistic-Full (x-axis) and Logistic-Reduced (y-axis) methods when the genotypes and covariates are moderately correlated. From left to right, the panels represent case-control ratios 1 : 1, 1 : 9 and 1 : 99, respectively. From top to bottom, the panels represent MAFs 0.001, 0.01 and 0.05, respectively.

**Figure 4:**
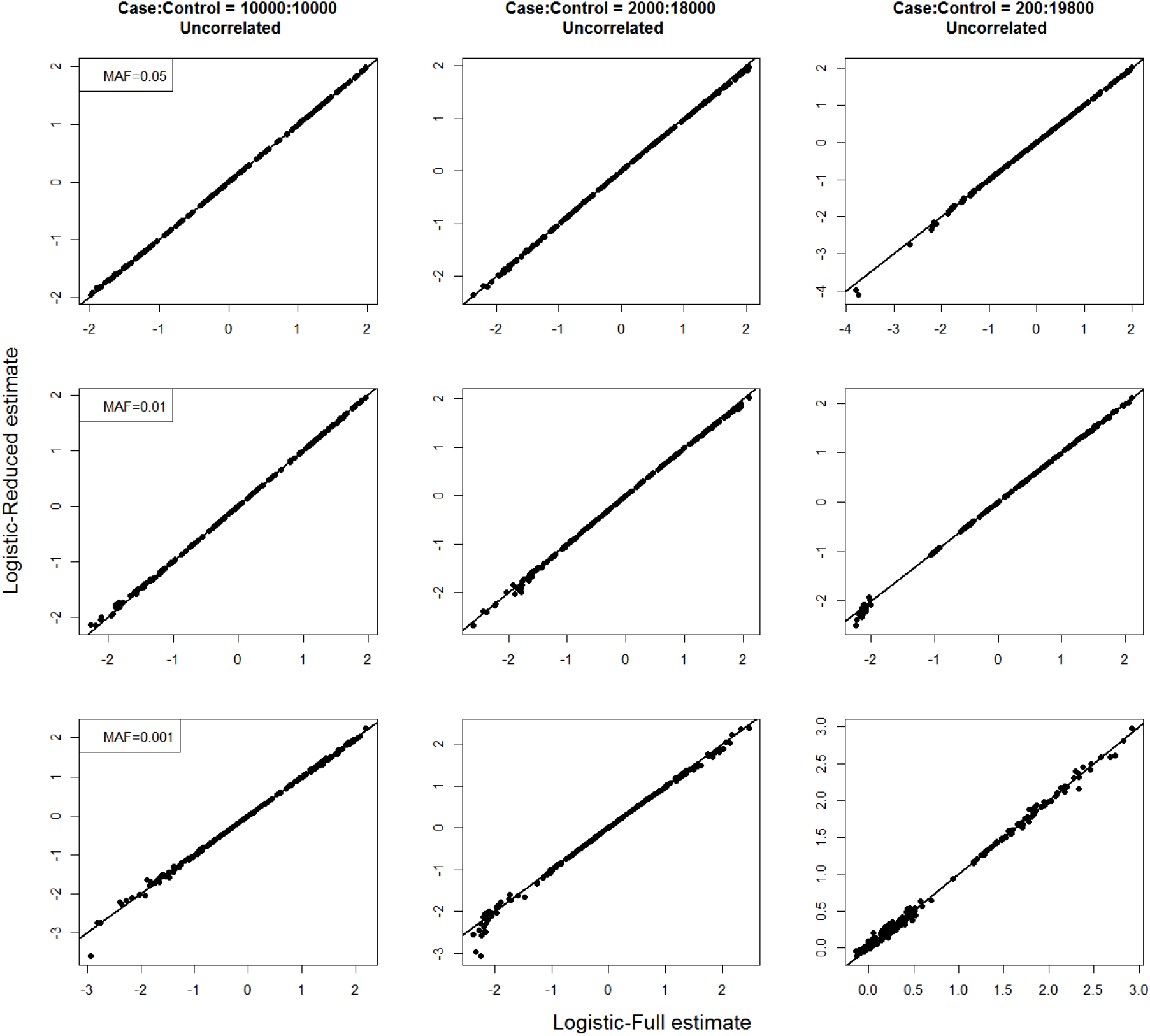
Scatter plots of estimated genotype LORs from Logistic-Full (x-axis) and Logistic-Reduced (y-axis) methods when the genotypes and covariates are uncorrelated. From left to right, the panels represent case-control ratios 1 : 1, 1 : 9 and 1 : 99, respectively. From top to bottom, the panels represent MAFs 0.001, 0.01 and 0.05, respectively.

**Figure 5:**
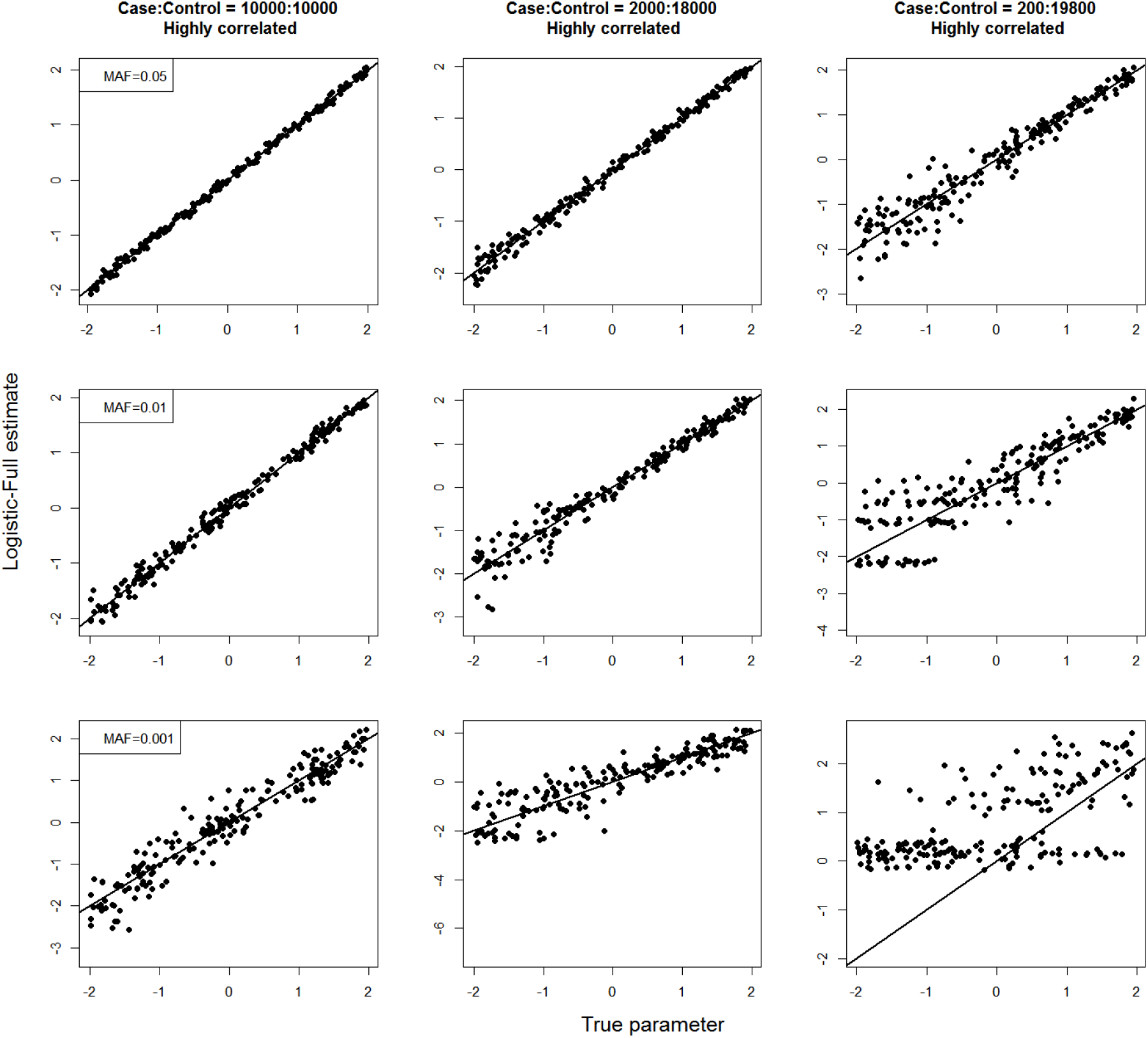
Scatter plots of the true parameter values (x-axis) and estimated genotype LORs from the Logistic-Full (y-axis) methods when the genotypes and covariates are highly correlated. From left to right, the panels represent case-control ratios 1 : 1, 1 : 9 and 1 : 99, respectively. From top to bottom, the panels represent MAFs 0.001, 0.01 and 0.05, respectively.

**Figure 6:**
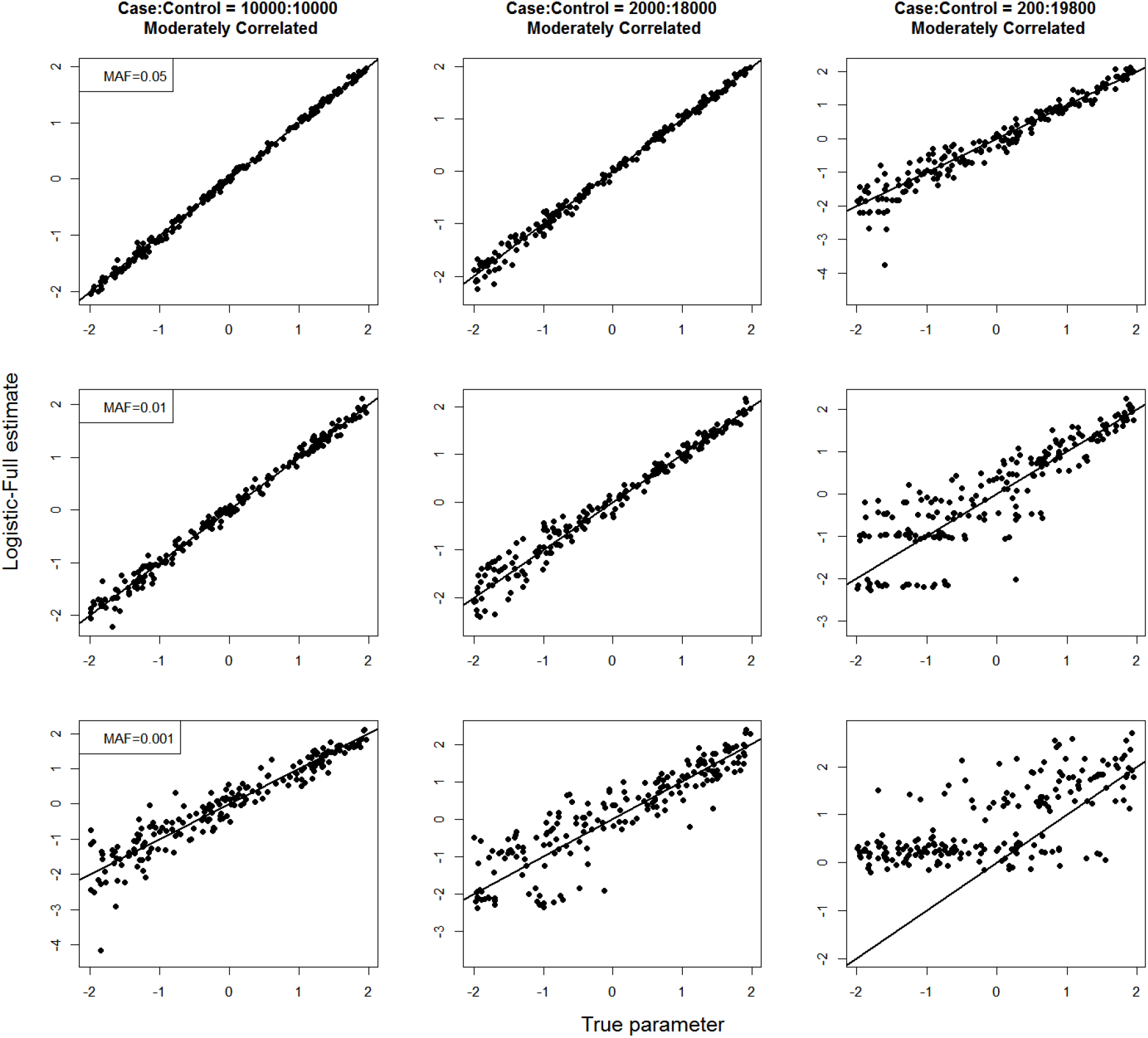
Scatter plots of the true parameter values (x-axis) and estimated genotype LORs from the Logistic-Full (y-axis) methods when the genotypes and covariates are moderately correlated. From left to right, the panels represent case-control ratios 1 : 1, 1 : 9 and 1 : 99, respectively. From top to bottom, the panels represent MAFs 0.001, 0.01 and 0.05, respectively.

**Figure 7:**
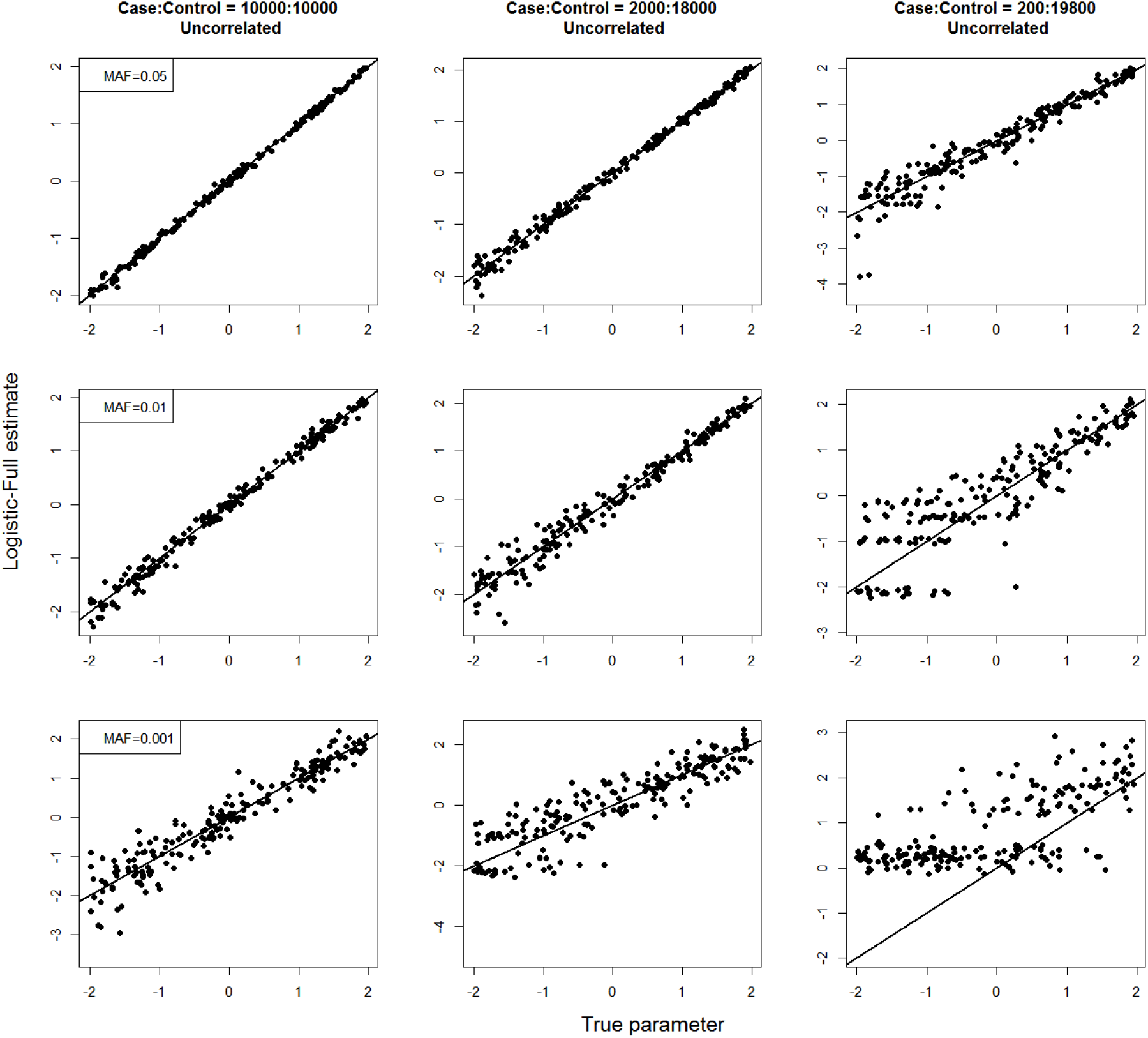
Scatter plots of the true parameter values (x-axis) and estimated genotype LORs from the Logistic-Full (y-axis) methods when the genotypes and covariates are uncorrelated. From left to right, the panels represent case-control ratios 1 : 1, 1 : 9 and 1 : 99, respectively. From top to bottom, the panels represent MAFs 0.001, 0.01 and 0.05, respectively.

**Figure 8:**
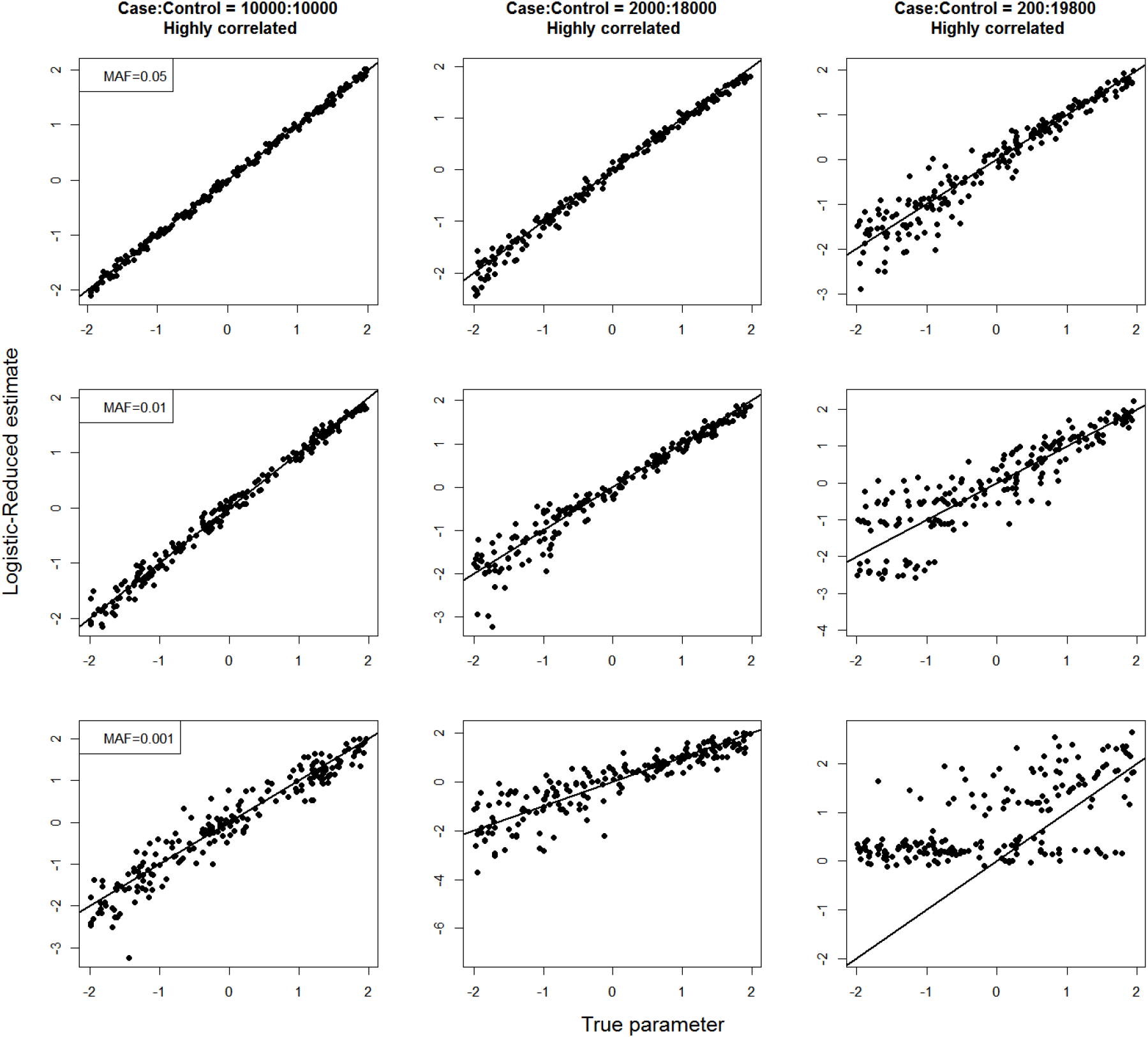
Scatter plots of the true parameter values (x-axis) and estimated genotype LORs from the Logistic-Reduced (y-axis) methods when the genotypes and covariates are highly correlated. From left to right, the panels represent case-control ratios 1 : 1, 1 : 9 and 1 : 99, respectively. From top to bottom, the panels represent MAFs 0.001, 0.01 and 0.05, respectively.

**Figure 9:**
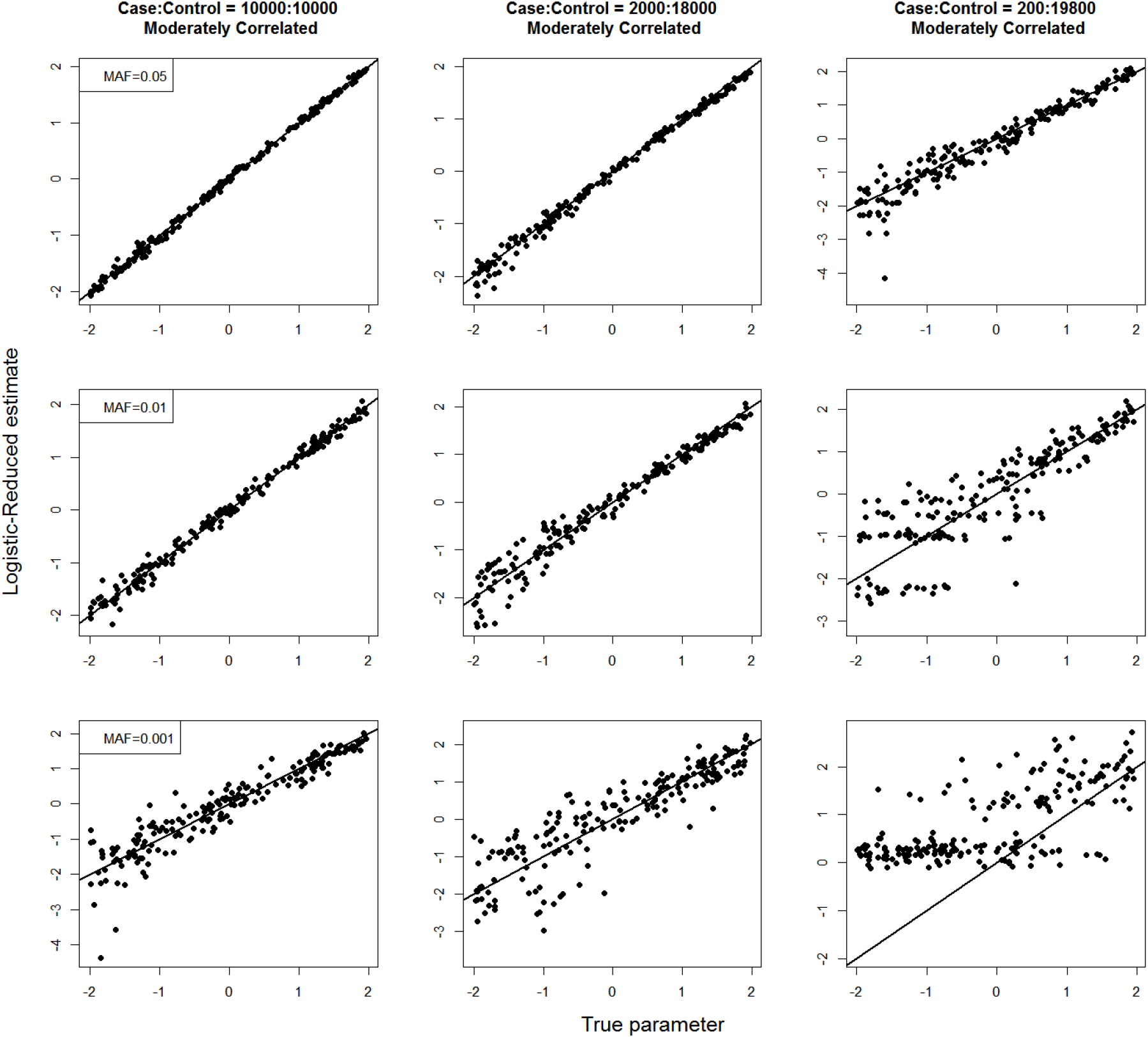
Scatter plots of the true parameter values (x-axis) and estimated genotype LORs from the Logistic-Reduced (y-axis) methods when the genotypes and covariates are moderately correlated. From left to right, the panels represent case-control ratios 1 : 1, 1 : 9 and 1 : 99, respectively. From top to bottom, the panels represent MAFs 0.001, 0.01 and 0.05, respectively.

**Figure 10:**
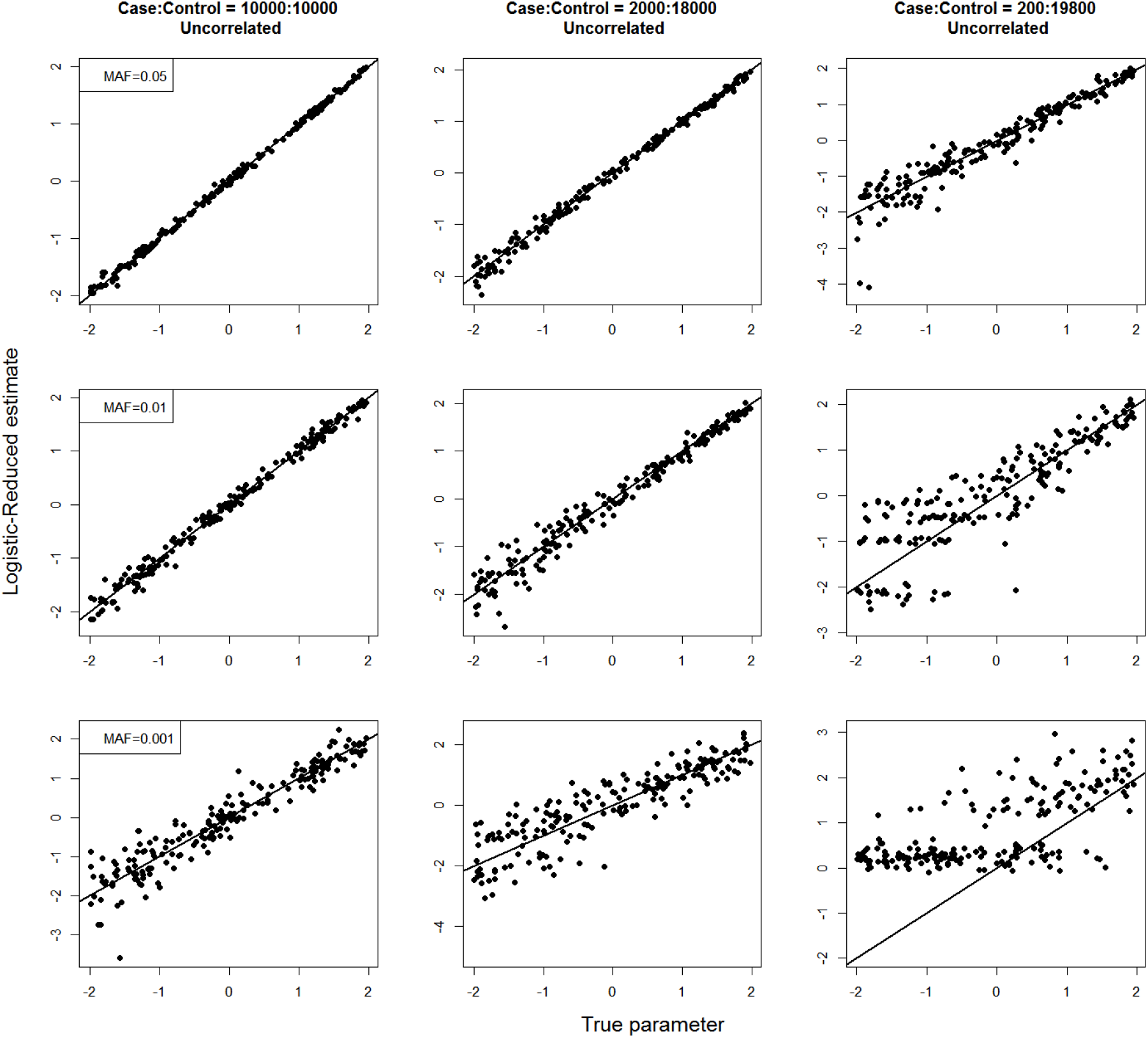
Scatter plots of the true parameter values (x-axis) and estimated genotype LORs from the Logistic-Reduced (y-axis) methods when the genotypes and covariates are uncorrelated. From left to right, the panels represent case-control ratios 1 : 1, 1 : 9 and 1 : 99, respectively. From top to bottom, the panels represent MAFs 0.001, 0.01 and 0.05, respectively.

Figure 1 presents the projected computation times for analyzing 10 million variants across 1500 phenotypes on a dataset of 20000 samples. The projected computation times clearly show that Logistic-Reduced is faster than Logistic-Full. For example, 197 CPU-years is required to analyze a moderately unbalanced dataset using Logistic-Full compared to only 16 CPU-years using Logistic-Reduced, when the number of covariates is *K* = 20. Moreover, the computation times of the Logistic-Full method increase at a faster than linear rate with the number of covariates, whereas the computation times of the Logistic-Reduced method remains almost the same.

### 4 Conclusion

We proposed a faster computation method for genotype LORs in case-control studies, which reduces the computation complexity from *O*(*nK*^2^ + *K*^3^) to *O*(*n*) compared to the traditional logistic regression with Firth bias-correction (Logistic-Full), where *n* is the sample-size, and *K* is the number of covariates. Our proposed method requires ~ 8–10x less computation time when analyzing a logistic regression model with 15 covariates, and ~ 16–20x less computation time when there are 25 covariates. Using simulation studies, we showed that for relatively smaller genotype LORs, the estimates from our method are almost identical to the estimates from the Logistic-Full method across different case-control ratios and rarity of alleles.

## Acknowledgments

The research was supported by NIH R01 HG008773.

